# Cultivation and genomic characterisation of novel methanogens from a desert biocrust

**DOI:** 10.1101/2025.09.22.677810

**Authors:** Weitao Tian, Eva Petrová, Sanae Sakai, Julius Eyiuche Nweze, Anne Daebeler, Roey Angel

**Author notes:** Institut National de Recherche Scientifique, Centre Armand Frappier Santé et Biotechnologie, Laval, QC, Canada. **Correspondence:** Roey Angel.

## Abstract

Methanogens are strictly anaerobic archaea capable of energy conservation by methane production, yet their presence in oxic and arid environments challenges existing paradigms. In this study, we enriched and genomically characterised seven methanogenic cultures from desert biocrusts, affiliated with the genera *Methanobacterium*, *Methanosarcina*, and *Methanocella*. Six of these new cultures represent new species. Nonetheless, phylogenomic analyses revealed close genetic relationships with organisms from anoxic environments, indicating the absence of an evolutionary distinction. Comparative genomics exposed diverse though non-unique repertories of antioxidant (e.g. catalase, superoxide dismutase and desulfoferrodoxin), and desiccation-resistance genes (including genes for maintaining osmotic pressure and repair of cell wall and membrane), with *Methanobacterium* spp. possessing the lowest gene abundance and diversity for oxygen and desiccation tolerance. Nevertheless, the occurrence of a Class I methanogen such as *Methanobacterium* in arid soils challenges the notion that members of this class are less oxygen tolerant than Class II. Pangenome analysis further uncovered unique genes enriched in membrane-associated functions and potentially non-functional stress-related genes. Via a global metagenomic survey, we find that methanogens are underdetected in dryland soils, likely due to sequencing depth limitations. Our findings highlight previously overlooked methanogen diversity and ecological plasticity in oxic and desiccated habitats, and emphasise the need for further studies to elucidate their survival strategies.

## 1 Introduction

Methanogens are a group of *Archaea* that reduce carbon dioxide (CO_2_), acetate (acetoclastic methanogenesis), or methyl compounds (methylotrophic methanogenesis) to produce methane (CH_4_) as a final metabolite [1, 2]. Methane has gained increasing attention for its significant contribution to climate change, as it is a 84-fold more potent a greenhouse gas than CO_2_ over a 20-year period [3]. Methanogens are the main biological source of CH_4_, and therefore play a crucial role in the global CH_4_ budget [4, 5]. They are strict anaerobes and are commonly found in anoxic environments, such as sediments, rice paddies, and animal guts [6, 7]. Methanogenesis, which predates the appearance of O in the atmosphere [8, 9], represents a form of anaerobic respiration [1, 10]. Phylogenetic analysis based on the SSU rRNA gene and conserved ribosomal protein markers showed methanogens are not monophyletic but comprise two distantly related groups, designated as Class I and Class II [11, 12]. Of the six recognised orders, Class I includes *Methanobacteriales*, *Methanococcales,* and *Methanopyrales*, and Class II includes *Methanosarcinales*, *Methanomicrobiales,* and *Methanocellales* [13–15]. Additionally, recent discoveries have further expanded the known methanogen diversity, including *Methanomassiliicoccales* [16, 17], and novel lineages in the phyla *Korarchaeota* [18], *Thermoproteota* (*Methanomethylicia*) [19], and unique methanogenic lineages outside *Euryarchaeota* [20].

In recent years, several methanogen genera have been detected in different environments that experience either a steady flux of oxygen, such as rice roots [15], wetlands [21], and lake water [22], or are mostly oxic, such as oxygenated sandy sediments, savannas, and even surface desert soils [23–27]. The genera *Methanobacterium*, *Methanosarcina* and *Methanocella* have consistently been found in the above-mentioned soils, especially in dryland areas [24, 28]. Moreover, methanogens in aerated soils have been shown to become active and grow when oxygen levels drop, for example after rewetting of desiccated soil and sediment ecosystems [22, 23, 28, 29]. The presence and activity of methanogens in desert soils is striking, as it implies that rewetting events may create transient anoxic microenvironments facilitating methane production, highlighting an overlooked source of atmospheric methane. These upland soil and desert methanogens endure both oxygen fluctuations and the osmotic pressures of desiccation and rehydration. Accordingly, laboratory experiments have confirmed that *Methanobacterium*, *Methanosarcina*, and *Methanocella* can survive in oxidative environments for hours or even days [24, 28, 30, 31]. However, unlike some other strict anaerobes (e.g. *Clostridium* sp. and *Sporomusa* sp.) [32], methanogens do not form spores, and the mechanisms they use to cope with oxygen and desiccation remain largely unknown.

Several key cofactors required for methanogenesis are sensitive to oxygen (O_2_) and its derivatives, i.e., reactive oxygen species (ROS), including hydrogen peroxide (H_2_O_2_), hydroxyl radicals (-OH), and O ^-^ radicals [30, 33–35]. In all the methanogenic pathways, the pivotal final step of the reaction of coenzyme B with the methyl group bound to the coenzyme M (CH_3_-S-CoM) to produce CH_4_ and the heterodisulfide (CoB-S-S-CoM) is highly conserved in methanogens and the most O_2_-sensitive [36]. The [4Fe-4S]^2+^ clusters contained in key enzymes involved in the process (heterodisulfide – HDR) and the low-coordinated iron atoms bound to the substrate make this step especially susceptible to ROS damage [37, 38]. Metagenomic analyses have unveiled a range of mechanisms by which methanogens may cope with oxidative stress, including synthesising antioxidant enzymes that detoxify ROS, reducing ROS production, and repairing DNA and protein damage [39–41]. Antioxidant enzymes rely on reducing equivalents, usually supplied by small redox proteins (including rubredoxin and thioredoxin) [33, 42]. Both classes of methanogens encode various antioxidant enzymes, but antioxidant-encoding genes are generally more abundant in Class II methanogens [43]. Additionally, methanogens possess repair enzymes such as MutS/MutL homologs [41], and methionine sulfoxide reductase [44], which protect against oxidative damage to nucleic acids and proteins, respectively. In response to desiccation stress, a variety of proteins in methanogens could play a protective role. Certain genera, including *Methanobacterium* and *Methanosarcina*, regulate osmotic pressure by accumulating compatible solutes, such as glycine betaine and proline, thereby maintaining cellular osmotic balance and stabilising proteins [45, 46]. Moreover, *Methanosarcina barkeri* has been shown to produce extracellular polymeric substances (EPS), which can help the cells retain more moisture and protect during desiccation [47]. *Methanosarcina* is also known to form biofilm that protect cells, with lipid synthesis upregulated to enhance membrane fluidity and aid in water retention [48]. Finally, homologues of late embryogenesis abundant (LEA) proteins, which are common in plants, have been identified in methanogens (e.g., *Methanocella paludicola*) [49, 50]. Such proteins may protect other proteins from being inactivated when partially dehydrated [49]. Despite these insights, most studies on methanogens in (transiently) oxic soils have focused on community-level changes in the environment, with limited molecular investigations regarding cellular mechanisms [28, 51]. Furthermore, all the studies using methanogen cultures outlined above were conducted using isolates from anoxic environments. This leaves a large gap in our understanding of the mechanisms responsible for ROS resistance in methanogens, which exist and survive in oxic conditions.

In this study, we enriched and cultured new methanogens from desert biocrusts, sequenced their genomes, and performed comparative genomics with 35 genomes and metagenome assembled genomes (MAGs) from multiple closely related lineages. Our study aimed to identify the genetic traits that enable methanogens inhabiting upland soils to adapt to aerated and desiccated conditions.

## 2 Materials and methods

### 2.1 Enrichment and cultivation, methane measurement, and growth assessment

Biocrust samples (ca. top 3 mm) were collected from two adjacent Negev Desert sites in Israel: near the ancient Nabatean city of Avdat (30°47′N, 34°46′E), and near the Liman irrigation system (30°50′N, 34°45′E). The area is an elevated, hilly region (ca. 800 m.a.s.l.) with an average annual rainfall of 80-100 mm, classified as arid [52]. The local loess soil is typically dry and exposed to air for most of the year [53]. Previous studies have shown the presence and activity of members of *Methanosarcina* and *Methanocella* in the biocrust, but not in the underlying soil [24].

To enrich methanogens, 1 g of biocrust was added to 5 ml of sterile deionised water in two 25 ml Balch-type tubes sealed with black butyl stoppers (VWR), crimped with an Al-ring, and flushed with an N_2_/CO_2_ (80%:20% vol/vol; see Supplementary Material for more details) gas mixture. Enrichment cultures were kept in the dark at 32 °C in a vinyl anaerobic chamber (Coy) without shaking. Methane production was measured weekly using a gas chromatograph (Agilent GC 6850) equipped with a ShinCarbon ST Micropacked GC column (Restek) and a Flame Ionisation Detector (FID, Agilent Technologies). After one month, 1 ml H_2_ (5% final conc.) and 1 ml of deoxygenated adapted Sekiguchi medium (pH 7.0, see Supplementary Material) were added [54]. After another month, CH_4_-producing enrichments were transferred (1:20) into fresh medium with N_2_/CO_2_ (80%:20%) headspace. Two enrichment lines were established: one for *Methanobacterium* with the addition of 5% H_2_ in the headspace, and another for *Methanosarcina* with Na-acetate (0.01 g l^-1^) in the medium. Methane production was monitored regularly. The community composition was monitored every few months via molecular methods as described below. After 10 months, 1 ml of *Methanobacterium*-dominated enrichment (34.4% of total community, 90.6% of total methanogens, estimated using ddPCR, see below) was transferred to 100 ml DSMZ 1523 medium [55] in 250 ml Duran bottles, capped with sterile butyl rubber stoppers, with 50% H_2_ added to the headspace. A *Methanosarcina-* dominated (91% of total community) culture was established after 16 months, for which a similar transfer was established into adapted DSM 960 medium ([56]; see Supplementary Material) with N_2_/CO_2_ (80%:20%) in the headspace.

To specifically enrich *Methanocella* sp., approximately 20 g of biocrust was pre-incubated at room temperature in a custom-made soil microcosm [28] under moist (field water-holding capacity of 9%) and anoxic (N_2_:CO_2_, 80%:20%) conditions for four weeks to activate the anaerobic microbial community. Afterwards, 5 g of pre-treated soil was mixed with 40 ml of fresh water basal medium (pH 7.0) [57] in a sterile 100 ml Duran bottle anoxically. 10% H_2_ and 5% sterile air were added to the headspace after flushing with N_2_/CO_2_ (80%:20%) and the culture was incubated anoxically at 37 °C. The addition of air was designed to suppress the growth of other methanogens that are assumed to be less aerotolerant than *Methanocella*. Methane production was monitored as stated above on a weekly basis. A mix of antibiotics was applied to all media to suppress bacterial growth. More details of media and enrichment lines are given in the Supplementary Material 1. Images of the enriched methanogenic strains were obtained using a scanning electron microscope (Supplementary Material 2).

### 2.2 DNA extraction, sequencing, and genome assembly

Soil biocrust DNA was extracted with the FastDNAä SPIN KIT for SOIL (MP Biomedicals), and culture DNA was extracted using a phenol-chloroform extraction method [58]. The proportion of methanogens to all prokaryotic community members in the cultures were determined through droplet digital PCR (ddPCR) on a Bio-Rad QX200 Droplet Digital PCR System using the following assays: universal prokaryotes (general 16S rRNA gene primers), universal methanogens methanogens (*mcrA* primers), and specific methanogen genera (genus-specific 16S rRNA primers) . Details of the primers used and the ddPCR assays are given in Table S1. The genus-specific ddPCR values were corrected to methanogen cell numbers by dividing by the number of 16S rRNA gene copies in the genomes of each target genus: 2 copies in *Methanobacterium* [59], 3 copies in *Methanosarcina* [60], and 2 copies in *Methanocella* [15].

When the cell numbers of *Methanobacterium*, *Methanosarcina*, and *Methanocella* reached of 1.9 ×10^6^, 2.1×10^6^, and 1.3×10^6^ per ml, respectively as determined by ddPCR, DNA was extracted from the cultures using the Quick DNA HMW Magbead Kit (Zymo Research) according to the manufacturer’s instruction after an overnight treatment of harvested cells in DNA/RNA shield solution (Zymo Research) at -20 °C. Total DNA was sent to Novogene Co., Ltd. (Munich, Germany) for library preparation and HiFi long-read sequencing on a PacBio Revio platform. The FASTQ sequences were extracted from the BAM file format using the ‘bam2fastq’ tool from the PacBio BAM toolkit [61]. The extracted reads were then *de novo* assembled with the ‘--meta’ command of the metaFlye algorithm (v2.9.3; [62]). The assembly outputs were visualised with the Bandage software (v0.8.1; [63]), and the contigs were exported in FASTA format. Circular genomes underwent further quality evaluation using QUAST (v5.2.0; [64]) to assess the assembly metrics, BUSCO (v5.7.0; [65]) to evaluate genome completeness based on conserved single copy orthologs, and CheckM (v1.2.2; [66]) to assess the level of contamination and completeness. Method details for DNA extraction and sequencing on an Illumina platform of an additional set of four cultures are given in Supplementary Material 3.

### 2.3 Phylogenomics and comparative genomics

PacBio- and Illumina-derived genomes were classified using GTDB-Tk (v1.4) with the archaeal reference database [67, 68]. To provide context, 35 reference genomes and MAGs (completeness > 80, contamination < 5) from methanogens of the same respective families (i.e. *Methanobacteriaceae*, *Methanosarcinaceae*, and *Methanocellaceae*) were obtained from the Genome Taxonomy Database. For each class, a set of 122 archaeal single-copy marker genes [68] was separately aligned, resulting in three alignment files [69–71] and trimmed using trimAl (v1.5; removed gaps > 50%) [72]. Afterwards, three maximum-likelihood trees were generated with IQ-TREE (v2.1.1) applying the WAG evolutionary model and with 1,000 bootstrap iterations [73]. One outgroup genome was included per tree (*Methanosarcina mazei* for *Methanobacterium* and *Methanocella*, and *Methanobacterium veterum* for Methanosarcina). The phylogenetic trees were visualised and annotated using iTOL [74]. Subsequently, the open reading frames (ORFs) of all genomes and MAGs were predicted using Prodigal [75]. A custom set of 60 HMM marker profiles (oxic and desiccation stress genes) was searched using ‘hmmsearch’ [76] with an e-value cutoff of 1.0×10^-10^. The antioxidant gene set was selected according to Lyu & Lu [41] and Johnson & Hug [99]. These included genes involved in combating oxidative stress (peroxides removal, free radical removal, iron storage / redox buffer, Fe-S cluster assembly, DNA glycolyase / endonuclease, and S=O / S-S recovery), and desiccation related damage (osmotic pressure regulation and membrane and cell wall repair) in methanogens. A complete list of all 61 marker proteins is given in Table S6. Methods for ordination analysis and a global search in publicly available metagenomes are given in Supplementary Material 4.

To allow species delineation, genome-wide Average Nucleotide Identity (ANI) and Average Amino Acid Identity (AAI) values were calculated between the 7 newly cultured strains and selected genomes of closely related methanogens for *Methanobacterium*, *Methanosarcina*, and *Methanocella*, according to [77, 78] respectively. The ANI and AAI heatmap matrices were visualised with ChiPlot [79].

### 2.4 Pangenome construction and annotation

To further compare the genomic composition of methanogens obtained in this study with closely related ones, we conducted a pangenome analysis using Anvi’o (v8; [80]). Unique and shared genes were identified as described in Eren, 2025 ([81]; detailed in Supplementary Material 5). Through the pangenome analysis, we identified distinct genes in each of the newly sequenced genomes (hereafter referred to as unique genes). These gene lists were manually explored for the presence of genes related to stress tolerance. Selected genes were quantified across all genomes presented in this study, and gene neighbourhoods of these stress-associated genes were visualised with the ‘Gene Graphic’ online tool [82].

## 3 Results and discussion

### 3.1 Cultivation and methanogenic activity

The occurrence of active methanogens in environments experiencing oxygen or periodic desiccation has been known for years [23, 24, 28, 31], but the genetic basis has not been systematically studied. In part, this is due to the lack of isolates and their genomes from such environments in public repositories. Here, we partially filled this gap by culturing and genomically characterising desert biocrust methanogens affiliated with genera systematically found in upland soils, namely with *Methanobacterium*, *Methanosarcina*, and *Methanocella*.

It is believed that cultures of methanogens require strictly anoxic conditions. These classical conditions, i.e. under H_2_/CO_2_ excluding O_2_, were applied in this study to successfully enrich *Methanobacterium* and *Methanosarcina*. However, members of the genus *Methanocella* are notoriously difficult to culture owing to their slow growth and reduced competitiveness with other H utilising methanogens [15, 85]. Leveraging the assumption that members of the *Methanocella* genus are the most oxygen-tolerant known methanogens [24], we regularly spiked the headspace of the *Methanocella* enrichment cultures with 5% lab air in order to favour them over the faster-growing *Methanosarcina* and *Methanobacterium* species.

The abundances of *Methanobacterium* spp., *Methanosarcina* spp., and *Methanocella* spp. in the original soil crust samples were 7.0×10^4^, 6.5×10^2^, and 1.3×10^2^ cells per gram dry weight, representing 2.4×10^-4^%, 2.6×10^-6^%, and 5.2×10^-7^% of the total prokaryotic communities, calculated and normalised from ddPCR results. Although abundances are extremely low, similar findings have been reported by Hall et al., who showed that aerotolerant methanogens present at low copy numbers in coastal sediments can nevertheless sustain ecologically significant methane emissions [23]. It suggests that rare methanogens, even when close to the detection limit, may still support critical ecosystem processes under favourable conditions [24, 29].

After eight, six, and two transfers of the *Methanobacterium*, *Methanosarcina*, and *Methanocella* cultures (i.e. on days 586, 503, and 129 of cultivation in each line), respectively, a strong dominance of each of the target methanogens was achieved. The enrichment levels of the target methanogen genus, expressed as a percentage of total methanogens, were 99.98% for *Methanobacterium*, 99.96% for *Methanosarcina*, and 98.05% for *Methanocella*, and of the entire prokaryotic communities were: 22.4% for *Methanobacterium,* 91% for *Methanosarcina*, and 16.3% for *Methanocella*. Methane production rates during enrichment cultivation were (1.13±1.10)×10^-3^, (7.37±7.75)×10^-3^, and (1.05±1.19)×10^-3^ nmol d^-1^ cell^-1^ for *Methanobacterium*, *Methanosarcina*, and *Methanocella* enrichment cultures, respectively. The methane production observed in these biocrust enrichments was 70-130 times higher than that reported for pure cultures of methanogens under controlled laboratory conditions [86–88]. This elevated activity may be attributed to the presence of coexisting microbial communities and residual organic matter, nutrients, and mineral components of the crust matrix.

The microscopic images obtained from scanning electron microscopy (SEM) analysis confirmed the typical cell morphologies and sizes for each methanogenic genus (Fig. 1).

**Figure 1.**
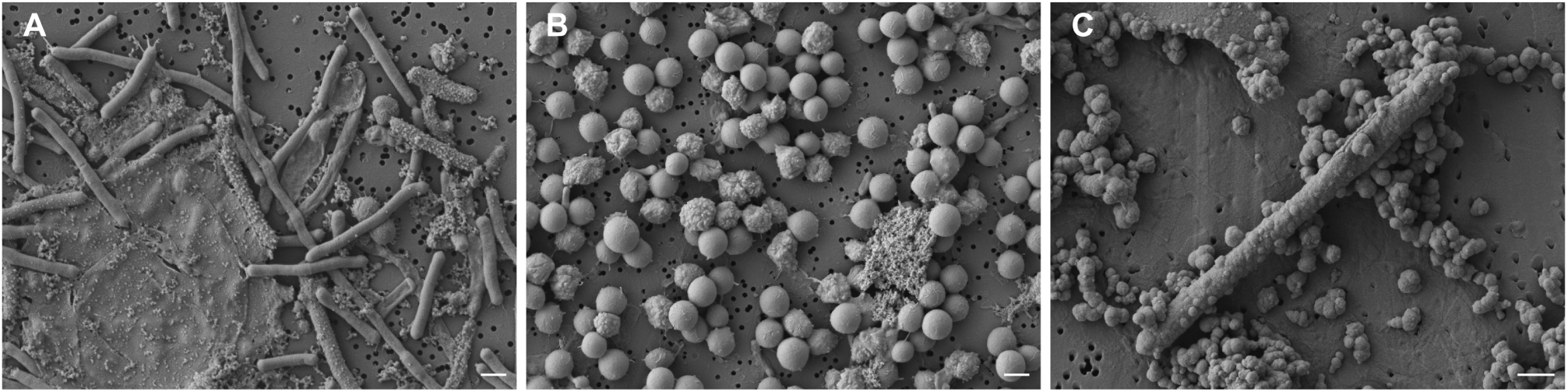
Scanning electron microscopy (SEM) images of the three newly cultured methanogens from desert biocrusts. *Methanobacterium* sp. (A) with a long rod shape, *Methanosarcina* sp. (B) with aggregated, coccoid cell clusters, and likely *Methanocella* sp. (C) with a shorter rod shape. The white line scale bars denote 1 µm.

Although the presence and even activity of methanogens in upland soils in general, and desert biocrust in particular, has been confirmed in the past [24, 28], the detection of Class I *Methanobacterium spp.* is particularly striking. This is because Class I are thought to possess a smaller repertoire of antioxidant enzymes [41], and more [4Fe-4S] clusters [40, 84], making them more vulnerable to transient oxidative conditions than Class II methanogens, such as *Methanosarcina* and *Methanocella* [40, 41]. Additionally, unlike Class II, Class I methanogens do not possess cytochromes, which allow for energy conservation via membrane-bound electron transport chains and fuel protective mechanisms linked to increased oxygen resistance [84]. The successful enrichment and cultivation of methanogens, including a cytochrome-lacking *Methanobacterium* from an environment that is mostly oxic, points to the possibility of the existence of yet-undescribed oxygen survival and detoxification strategies in these organisms.

These new enrichment cultures provide a valuable addition to the known methanogen cultures, which have all been obtained from anoxic habitats. They highlight a wider-than-expected ecological plasticity of methanogens and present a unique tool to further investigate potential oxygen tolerance and detoxification mechanisms.

### 3.2 Genome characteristics and phylogeny

Using *de novo* assembly, we obtained seven distinct high-quality methanogen genomes from our enrichment cultures, four of which were circular: three classified as *Methanobacterium* spp. and three as *Methanosarcina* spp. (each two MAGs from the Illumina NovaSeq sequencing and one closed genome from PacBio sequencing), as well as one closed genome classified as belonging to a *Methanocella* sp. (closed genomes are shown in Fig. S1). Quality assessment using QUAST showed genome completeness of non-circular MAGs were 92.8, 93.6 (*Methanobacterium* spp.), and 99.7, 99.8% (*Methano*sarcina spp.), with minimal contamination levels evaluated by CheckM at 0, 4 (*Methanobacterium* spp.), and 0.98, 0 (*Methano*sarcina spp.), respectively. Likewise, minimal contamination levels of closed genomes were determined with 0.8, 0.65, and 0% for the *Methanobacterium* sp., *Methanosarcina* sp., and *Methanocella* sp., respectively.

Phylogenomic analysis confirmed the affiliation of the organisms in our enrichment cultures with the *Methanobacteriales*, *Methanosarcinales*, and *Methanocellales* orders (Fig. 2, Table S2). Unexpectedly, the new methanogenic cultures from a desert environment did not fall into separate clusters but showed to be closely related to several known taxa from generally anoxic habitats such as wastewater, groundwater, permafrost, sub-sea sediments, or rice paddies. Moreover, we did not detect any phylogenomic clustering correlating with the environmental conditions in which the organisms live. The average genome sizes and G+C contents across the examined *Methanobacterium*, *Methanosarcina*, and *Methanocella* genera were 2.62 Mb and 35.9%, 4.39 Mb and 40.9%, 2.39 Mb and 50.4%, respectively. The variations in genome size and GC content among these methanogens may reflect their distinct metabolic strategies and environmental niches. Commonly, a lower GC content, such as found for the *Methanobacterium* spp., is associated with adaptation to anaerobic conditions [89, 90], which aligns with the classical view on Class I methanogens.

**Figure 2.**
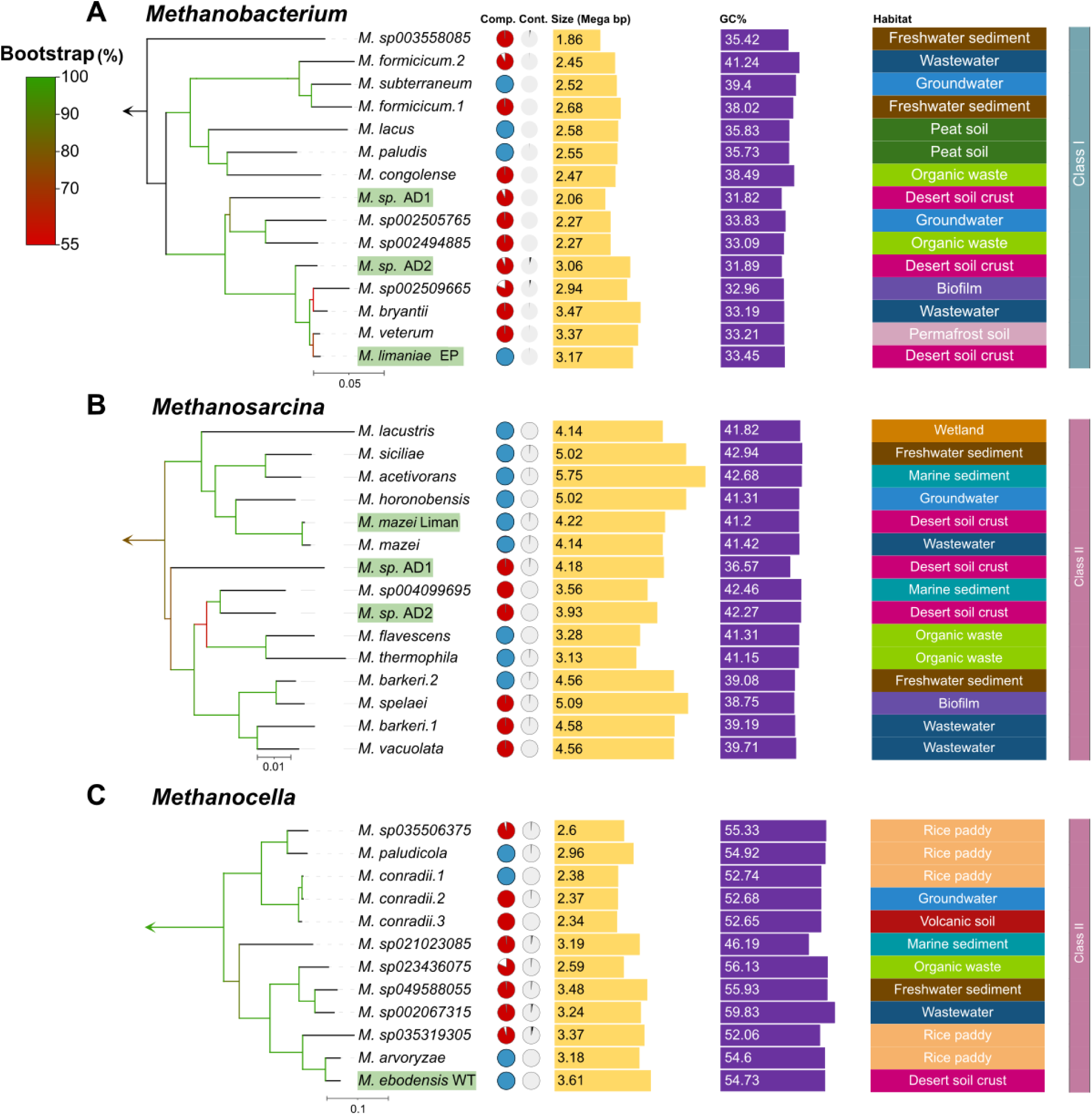
Phylogenomics and basic genome-assembly properties of the newly cultured methanogens and known representatives. Genome-based, maximum-likelihood trees for the genera *Methanobacterium* (a), *Methanosarcina* (b), and *Methanocella* (c) are shown with the newly cultured methanogens of this study highlighted in green. Colours on branches show bootstrap support value (%) on 1,000 replicates. The scale bars denote 0.05, 0.01, and 0.1 estimated substitutions per amino acid, respectively. The arrow at the base of each tree denotes rooting based on an outgroup (see Materials and Methods for specifications). Comp., Completeness (in red; blue solid circles represent circular genomes); Cont., Contamination.

### 3.3 Species delineation and distribution in global oxic drylands

Genome-wide average nucleotide identity (gANI) and average amino acid identity (AAI) based analyses were performed on the same *Methanobacterium*, *Methanosarcina*, and *Methanocella* datasets to classify the newly cultured desert methanogens on the species and genus levels and higher taxonomic ranks (Fig. 3). Based on the gANI analysis and a species level threshold of 96.5% [91] the three newly cultured *Methanobacterium* organisms represent separate species from each other and from *Methanobacterium veterum* and *Methanobacterium bryantii* (Fig. 3, Table S3) as their closest cultured relatives. For the species from which we obtained a closed genome, we therefore tentatively propose the name “*Candidatus* Methanobacterium limaniae EP”. We refrain from proposing new species names for the other two cultures, since they were only transiently present in the enrichment culture and we did not obtain circular genomes from them. Henceforth, we refer to these organisms as *Methanobacterium* sp. AD1 and AD2. Likewise, two of the newly cultured *Methanosarcina* organisms are separate species from each other and from *Methanosarcina flavescens* and *Methanosarcina barkeri* 2, as their closest cultured relatives. We refer to these organisms as *Methanosarcina* sp. AD1 and AD2 from hereafter. The third *Methanosarcina* organism was not a separate species from *Methanosarcina mazei,* and we therefore give the strain name “Liman” to this organism. Finally, the newly cultured *Methanocella* organism was shown to represent a new species, distinct from its closest cultured relative, *Methanocella arvoryzae*. We therefore tentatively propose the name “*Candidatus* Methanocella ebodanensis WT”. In summary, we have cultured and identified six novel methanogen species through a genomic analysis of enriched organisms. The difficulty in cultivating and isolating methanogens, especially those with specialised metabolic requirements, has long been a challenge in microbial ecology. The identification of new species from three genera inhabiting oxic desert biocrusts in this study highlights the underappreciated diversity of methanogens in oxic habitats.

**Figure 3.**
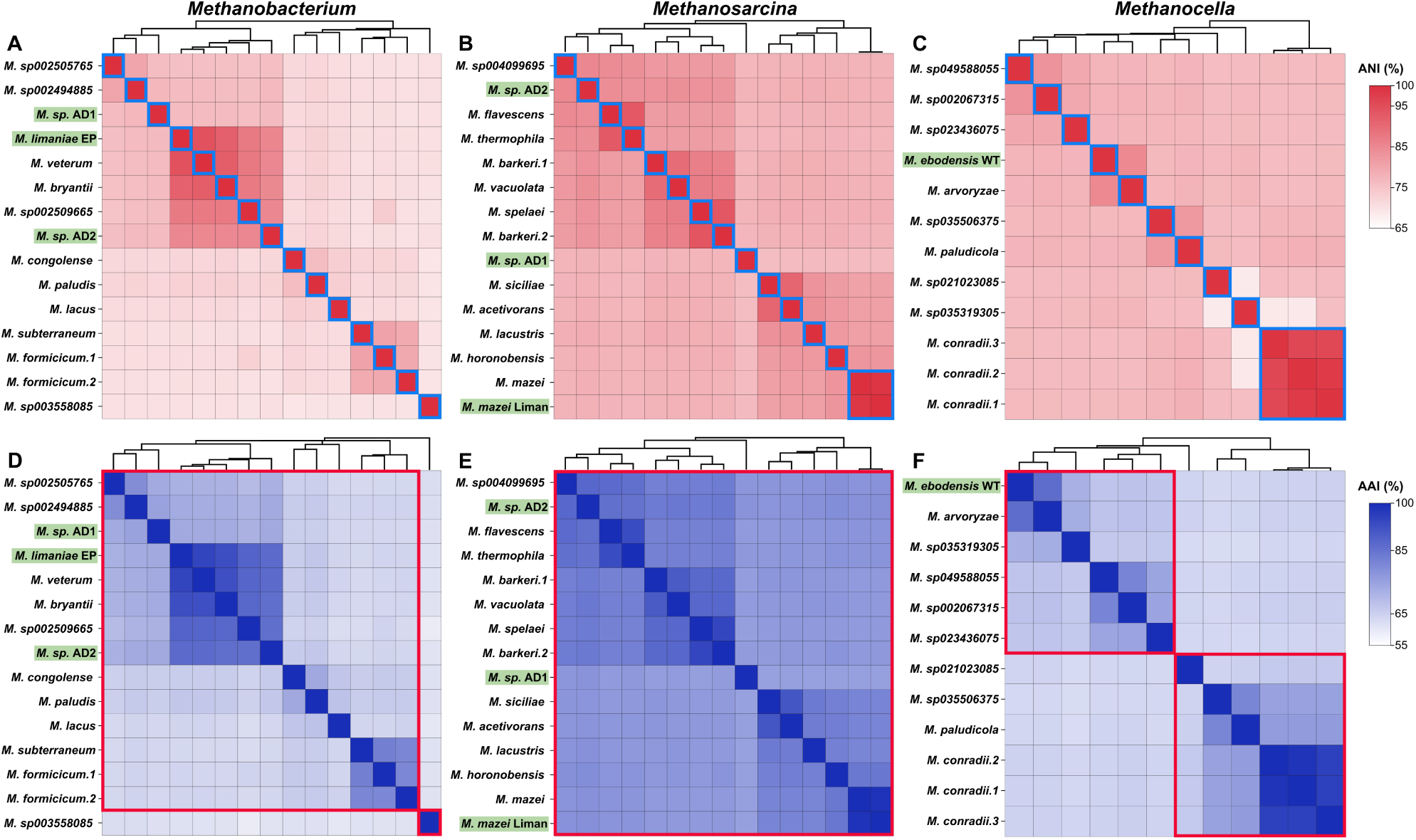
Genome-wide average nucleotide identity (gANI) and average amino acid identity (AAI) heatmaps for *Methanobacterium* (A, D), *Methanosarcina* (B, E), and for *Methanocella* (C, F). Names of genomes from this study are highlighted in green. Genomes within the same species in ANI heatmaps are framed in blue boxes, and genomes belonging to the same genera in AAI heatmaps are framed in red boxes.

To assess the distribution of methanogens in arid oxic soils worldwide, we surveyed 24 metagenomic datasets collected from the NCBI Bioproject database generated from samples from Europe, the Americas, Asia, Africa, Oceania, and Antarctica. Methanogens were detected in only five of the 24 projects (44 datasets, Table S4, S5). Among the detected methanogens, members of the class *Methanosarcinia* were most frequently observed. In addition, members of other methanogen classes, including *Methanomicrobia*, *Methanococci, Methanobacteria*, and *Methanopyri,* were all present in more than one dataset. However, the relative abundances were consistently low, representing max. 0.28% of the total microbial communities. Considering the discrepancy of apparent methanogen absence as assessed by metagenomic sequencing with the proven presence of methanogens via targeted approaches in this and previous work (e.g. for Negev desert biocrust [28]), we assume that the low detection rate of the global search reflects insufficient sequencing depth rather than their absence from these systems. These results highlight the fact that metagenomic surveys still overlook many rare, yet ecologically important taxa.

### 3.4 Genomic potential for antioxidant and desiccation-resistance mechanisms in the newly cultured methanogens

To identify genes associated with oxidative stress resistance and desiccation tolerance in the newly cultured methanogens, we conducted a targeted comparative genomic analysis across the genomes of our new cultures and those of their closely related relatives. There was a clear distinction in the encoded diversity of oxygen and desiccation stress genes between the three investigated genera, indicating their different potential stress response strategies (Fig. 4, Fig. S2, Table S7). Notably, *Methanosarcina* genomes exhibited higher completeness in Fe-S cluster assembly and DNA glycosylase / endonuclease genes (Fig. 4B), which may confer an adaptive advantage to maintain genomic integrity even under harmful conditions that cause DNA lesions. Interestingly, the antioxidant gene set in *Methanocella* species was not the most complete among the three genera (Fig. 4C). This finding was unexpected, considering the generally attributed highest relative tolerance to oxygen of *Methanocella* and their ecological niche as rice-root methanogens [92, 93].

**Figure 4.**
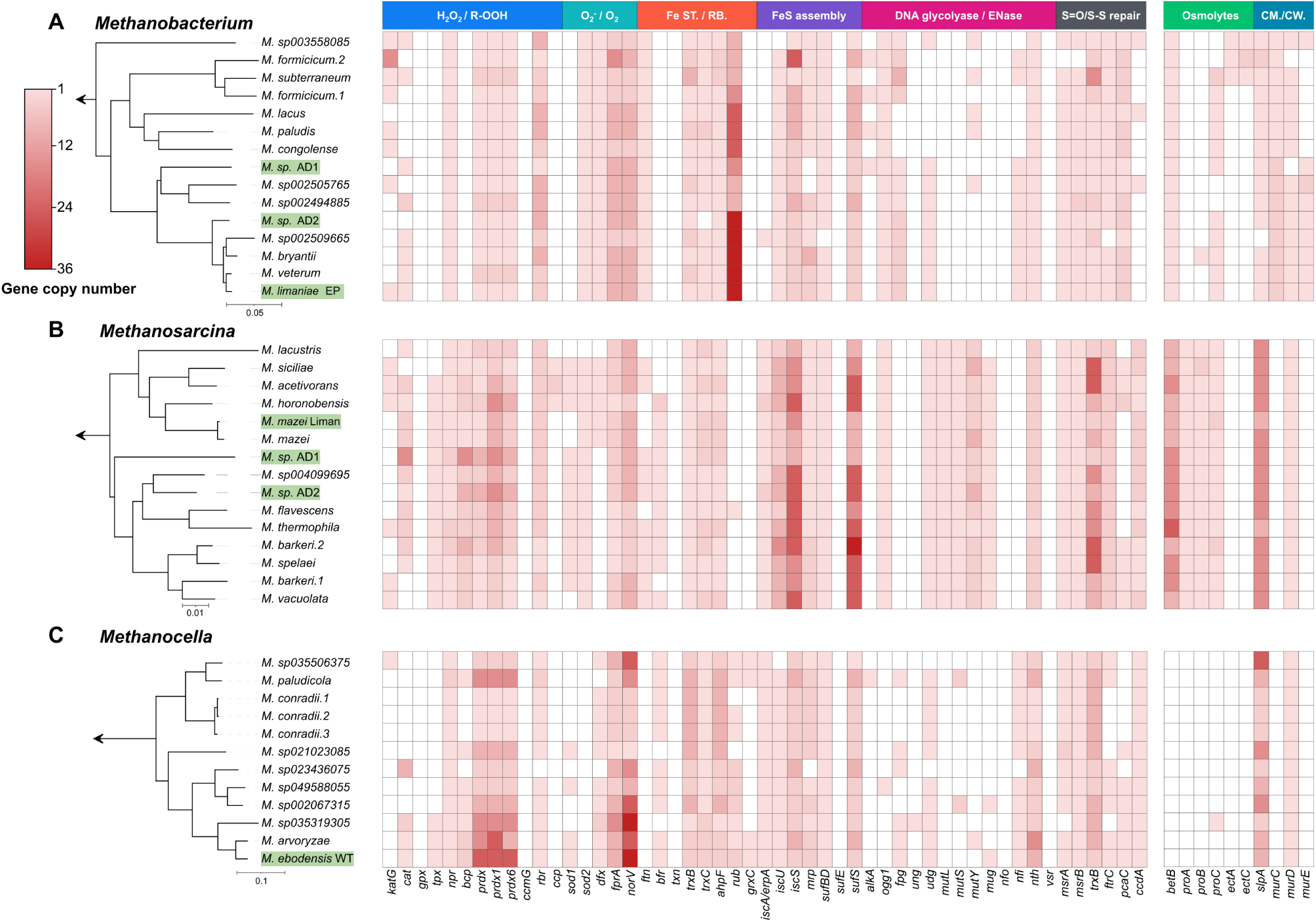
Distribution of distinctive genes related to antioxidant and desiccation stress traits in methanogens of the *Methanobacterium* (A), *Methanosarcina* (B), and *Methanocella* (C) genera. Genomes used in the analysis and phylogenomic trees on the left correspond to those in Fig. 1. A homology-based search for functional genes was performed using HMMER’s hmmsearch and manual examination. Solid and open squares indicate the presence and absence of the genes, respectively. The presence of multiple copies is indicated by colour intensity. Gene names are given at the bottom (see Table S6 for full names), and the corresponding functions are annotated at the top. Fe ST., iron storage; RB., redox buffer; CM./CW., cell membrane/cell wall.

Surprisingly, the enriched methanogens from this study did not show a higher diversity, higher gene copy numbers, or specific patterns compared to their sister taxa within the genera. However, “*Ca.* Methanocella ebodanensis WT” showed a modest increase in average copy number of antioxidant genes, ranking highest among the *Methanocella* genomes in this study. It aligns with our cultivation experiment, in which *Methanocella* demonstrated comparatively greater tolerance to O_2_ exposure than members of the *Methanobacterium*. The overall trends on the genus level remained apparent in our newly sequenced genomes, with the three *Methanosarcina* genomes containing the highest diversity and copy numbers and the *Methanobacterium* genome containing the least (Fig. 4, Table S7). These findings likely reflect long-term evolutionary adaptations rather than short-term responses to environmental pressures, emphasising that genomic potential for stress tolerance in methanogens is likely more constrained by phylogenetic lineages than by the immediate habitat.

Additionally, we specifically investigated the presence of LEA protein homologs (late embryogenesis abundant proteins) that were previously reported in Methanocella paludicola and annotated as such by Campos et al. [49]. In plants, LEA proteins protect against dehydration stress by stabilising proteins and membranes [94, 95], and have been discussed to provide similar functions in methanogens. However, our search using multiple archaeal LEA protein sequences from UniProt found zero hits across all Methanocella genomes, including M. paludicola. To further examine this discrepancy, we compiled 19 LEA protein sequences from plants, bacteria, archaea, and the five Methanocella proteins annotated by Campos et al. as LEA proteins, and constructed a neighbour-joining phylogenetic tree (Fig. S3). In this tree, the five putative LEA proteins from Methanocella clustered separately. Finally, a BLASTx search on the NCBI server showed that the putative LEA proteins from Methanocella have a high sequence similarity with KGG domain containing proteins and general stress proteins rather than with LEA proteins. These results raise doubts about whether methanogens, including Methanocella, truly harbour LEA proteins.

### 3.5 Pangenomic analysis and potentially non-functional genes

To further explore genomic differences between the newly cultured desert methanogens and their close relatives, we conducted a pangenome analysis. The analysis identified both core genes shared across all genomes and unique genes specific to single genomes. All seven genomes of our newly cultivated methanogens contained unique genes, with the genome of “*Ca.* Methanocella ebodanensis WT” containing the largest set (Fig. S4, Table S9). Strikingly, many (16%) of these unique genes in all our genomes were associated with membrane and cell wall biogenesis. In comparison, such genes accounted for 15% in *Methanobacterium*, 7% in *Methanosarcina*, and 10% in *Methanocella* among published genomes included in this study. The relative enrichment of such genes in our desert-derived strains may indicate lineage-specific strategies for maintaining cell integrity in the multi-factor extreme environment.

Additionally, several functionally intriguing genes were found, including those annotated as encoding subunits of glycosyltransferase (*rfaB*, *wcaA*, and *wcaE*) and the nitrous oxide reductase subunit D (*nosD)*. Glycosyltransferases are typically involved in exopolysaccharide biosynthesis and cell envelope modification, processes that may support stress resistance in bacteria [96, 97], but are poorly characterised in methanogens. The *nosD* gene encodes an accessory protein associated with the Nos operon for N_2_O reduction in denitrifiers [98], yet methanogens are not known to perform denitrification. To assess the distribution of these four genes, we quantified their copy numbers across all selected genomes in this study based on the anvi’o annotation (Fig. S5). The glycosyltransferase genes were widespread without genus-specific patterns. In contrast, *nosD* was enriched in *Methanosarcina* genomes only, implying a genus-specific functional adaptation possibly related to protein maturation. However, the putative *nosD* genes were not enriched in the genomes of the new desert methanogens, pointing towards a more general role.

To further understand the potential function of the gene annotated as *nosD* in our newly cultured methanogens, we performed a BLAST search against the NCBI nr protein database and out of the seven genes annotated as NosD, only three gave hits designated as NosD, albeit unverified (Table S8). A Neighbour-joining phylogenetic tree built with *bona fide* NosD sequences showed that our putative NosD protein sequences formed a distinct cluster, separate from the other known ones (Fig. S6). Additionally, synteny analysis of the putative *nosD* genes showed that other *nos* operon genes were absent (Fig. S7), suggesting that these genes may have alternative, yet unknown functions.

In summary, although certain genes with intriguing annotations such as *nosD* and glycosyltransferases were found among the unique genes, their unclear functional roles, lack of operon context, and phylogenetic divergence suggest that they may represent non-functional remnants or genes with alternative, yet-to-be-elucidated roles in methanogens.

## Conclusions and perspective

The occurrence of methanogens of the *Methanobacterium*, *Methanosarcina*, and *Methanocella* genera in many oxic and dry soils defied basic assumptions about their physiology and left open questions regarding the restricted occurrence of only these genera and their genomic machinery to persist under the stressful conditions of these habitats. Our characterisation of newly cultured methanogens showed that while they are equipped, to some extent, to handle ROS and desiccation, their genomes reveal no unique mechanisms that would set them apart from their relatives living in anoxic and wet environments. Moreover, we found that *Methanocella*, which is regarded as the most oxygen tolerant methanogen, is in fact depleted in many known antioxidant genes compared to its sister genera. Concomitantly, we found that *Methanobacterium*, which was predicted to be particularly oxygen sensitive, is not only prevalent in oxic soils but also well-equipped to handle oxygen and desiccation stressors. Given the lack of strong genomic signals in the newly cultured methanogens from desert biocrust, it is possible that differences in gene expression contribute to their apparent stress tolerance. Therefore, besides ongoing cultivation and genomic sequencing attempts, further work on gene expression under physiologically relevant stress conditions is required to elucidate the survival mechanisms of methanogens living in oxic and dry environments.

### Taxonomic considerations of “*Candidatus* Methanobacterium limaniae” sp. nov

Li.ma’nae. N.L. gen. n. limaniae, belonging to the Liman, from Greek *λιμήν* through Turkish and Russian, originally meaning a bay or a port, but in this context refers to a constructed runoff catchment, from which the strain has been isolated.

Phylogenetically affiliated with the genus *Methanobacterium*, phylum *Methanobacteriota*. The genome comprises one circular chromosome of 3,168,381 bp. The DNA G + C content is 33.45 mol%. Strain “*Ca.* Methanobacterium limaniae EP” was cultivated from biocrust of the Negev desert, Israel. Soil hydrogenotrophic methanogen with a rod-shaped morphology. Cells are slender and elongated, typically measuring ∼0.5 µm in diameter and 4-5 µm in length. The strain was routinely cultured with 80% H_2_ at 32°C in DSM17711 medium.

### Taxonomic considerations of “*Candidatus* Methanocella ebodensis” sp. nov

E.bo.den’sis. N.L. fem. adj. ebodensis, named after Eboda, a significant ancient Nabataean city along the “Incense Route”, close to the location where the strain was isolated. The Nabataeans were renowned for their sophisticated water collection techniques, which enabled them to thrive in the desert environment.

Phylogenetically affiliated with the genus *Methanocella*, phylum *Halobacteriota*. The genome comprises one circular chromosome of 3,614,771 bp. The DNA G + C content is 54.73 mol%. Strain “*Ca.* Methanocella ebodanensis WT” was cultivated from biocrust of the Negev desert, Israel. Soil hydrogenotrophic methanogen with a rod-shaped morphology. Cells typically measure ∼0.8 µm in diameter and 7-8 µm in length. The strain was routinely cultured with 10% H_2_ at 37°C in fresh water basal medium and subjected to 5% air regularly.

## Data availability

All sequencing data generated in this study have been deposited in the NCBI Sequence Read Archive (SRA) under BioProject accession number PRJNA1242298. Complete genome assemblies are available in NCBI GenBank under accession numbers CP187259.1, CP187260.1, and CP187261.1, and draft genome assemblies are available under accession numbers JBNAKP000000000.1, JBNAKN000000000.1, JBNAKM000000000.1, and JBNAKO000000000.1. The names of the novel methanogen strains “*Candidatus* Methanobacterium limaniae” sp. nov and “*Candidatus* Methanocella ebodensis” sp. nov have been validly registered under the SeqCode with the registry accession seqco.de/r:lhm19uk7 (https://seqco.de/r:lhm19uk7).

## CRediT author contribution

**WT, EP, AD and RA:** conceptualisation, data curation, methodology; **RA:** funding acquisition; **EP, RA:** project administration, resources; **WT, JEN:** formal analysis, software, visualisation; **WT, EP, SS:** investigation; **EP, AD, RA, SS:** supervision, validation; **WT, AD, RA:** original draft; **WT, EP, SS, JEN, AD, RA:** writing – review & editing.

## Acknowledgments

WT, EP, AD and RA were supported by the Czech Science Foundation (Standard Grant no. 22-34650S). Electron microscopy images were obtained thanks to the help of the BC CAS core facility Laboratory of Electron Microscopy supported by the Czech-BioImaging large RI project (LM2023050and OP VVV CZ.02.1.01/0.0/0.0/18_046/0016045 funded by MEYS CR). We thank Prof. Osnat Gillor from Ben-Gurion University of the Negev, Israel, for providing us the soil samples from which the methanogenic strains were enriched. We thank Ronnie Hirsch for her help in constructing the Latin nomenclature of the new species.

