## Supplemental methods for "Cultivation and genomic characterisation of novel methanogens from a desert biocrust"

**Supplementary Materials**

**1. Media and enrichment lines**

The adapted Sekiguchi medium contained the following components per litre: 0.3 g L-Cysteine monohydrochloride, 0.15 g KH_2_PO_4_, 0.15 g CaCl_2_·2H_2_O, 0.2 g MgCl_2_·6H_2_O, 0.5 g NH_4_Cl, 2.5 g NaHCO_3_, 1 ml resazurin solution (25 mg l^-1^), 1.5 ml Na_2_S (25g l^-1^), 3 ml coenzyme M (2 g l^-1^), 10 ml vitamin solution (according to DSMZ 141 [1]), 10 ml trace elements solution (according to medium DSMZ 318 [2]). Vitamins, trace elements, coenzyme M, resazurin, and Na_2_S were added from sterile anoxic stock solutions through a 0.2 µm SFCA filter (Thermo Scientific) using a syringe after autoclaving the medium. The atmosphere in incubation vessels was exchanged to N_2_:CO_2_ (80%:20%; 5.0 grade; Messer) using a self-constructed gassing manifold equipped with an oil vacuum pump that alternates between -0.9 and +0.9 atm. relative to the atmosphere, for 10 min. Cultures were routinely transferred by diluting (1:100) into fresh medium roughly every two months. To limit the growth of bacterial cells in the cultures, a mixture of the following antibiotics was applied to each transfer: kanamycin (MP Biomedicals), ampicillin, penicillin G (Merck), and vancomycin (Carl Roth) each at 50 mg ml^-1^.

The methanogen-dominated enrichment lines were maintained in strain-specific media and culture conditions as described below. One ml of *Methanobacterium*-dominated enrichment line was transferred to 100 ml medium (DSMZ 1523 [3]) in 250 ml Duran bottles, capped with sterile bromobutyl rubber stoppers with 50% H_2_ added to the headspace. The *Methanosarcina*-dominated culture was transferred to DSM 960 medium (originally designed for *Methanocella paludicola* DSM 17711 [4]) but supplemented with 1% methanol, 5 g l^-1^ NaCl, 0.3 mg l^-1^ FeSO_4_ to maintain and increase biomass. Antibiotics were applied to all media as described above.

For the *Methanocella* enrichment line, the medium was adapted from freshwater basal medium [5] and contained (per litre): 0.535 g NH_4_Cl, 0.136 g KH_2_PO_4_, 0.204 g MgCl_2_, 0.147 g CaCl_2_·2H_2_O, 2.52 g NaHCO_3_, 0.2 ml resazurin solution (25 mg l^-1^), 1 ml vitamin solution (as referenced above), 1 ml trace elements solution (as referenced above), 1 ml Selenium/Tungsten solution (according to [6]), and 1 ml penicillin (50 mg ml^-1^) and kanamycin (50 mg ml^-1^) antibiotics mixture solution. All components except vitamins and antibiotics were added to 1 l water. The headspace was evacuated and flushed with N_2_/CO_2_ (80/20%) for 10 min before autoclaving. Subsequently, the other components and a reducing agent (Na_2_S, 0.1 g l-1) were added through a 0.2 µm SFCA filter using a syringe.

2. Scanning electron microscopy

To prepare the cells for electron microscopy, 1 ml of culture was fixed in 2.5% glutaraldehyde (Carl Roth) in the respective medium for 3.5 h in an anaerobic chamber (Coy). The fixed cells were then filtered on to a 0.20 mm polycarbonate filter (Whatman), washed with diluted phosphate buffer saline (1:2 in PCR-grade water, pH 7.2), and post-fixed in 1% osmium tetroxide (Carl Roth) in PBS for 45 min. Afterwards, the cells were washed again with diluted phosphate buffer saline, and serially dehydrated in ethanol solutions (each 3 min in 25, 50, 70, 80, 90 in PCR-grade water and in 100%). Filters with cells were then sputter-coated with gold (Leica EM ACE200; Leica Microsystems) prior to imaging with a JSM-7401F scanning electron microscope (JEOL), operated at a 4 kV acceleration voltage.

For *Methanobacterium* enrichment culture used for SEM, methanogens accounted for 70.2% of the total prokaryotes, with Methanobacterium representing 76.5% of the methanogens. For Methanosarcina enrichment culture, Methanosarcina comprised more than 99.9% of both prokaryotes and methanogens. And for Methanocella enrichment culture, methanogens accounted for 84.2% of the total prokaryotes, consisting exclusively of *Methanosarcina* and *Methanocella* at approximately a 1:1 ratio. All values were derived from normalized ddPCR results.

3. DNA extraction, Illumina sequencing, and genome assembly

Four samples from *Methanobacterium* and *Methanosarcina* enrichment lines originating from Avdat biocrusts (each two samples) were sequenced using a Illumina HiSeq 4000 platform (Novogene GmbH) to generate paired-end reads. DNA was extracted with a phenol-chloroform method [7], purified using ethanol precipitation and quantified using PicoGreen dsDNA (Thermo Scientific) and sent for sequencing at Novogene Co., Ltd. Adapter sequences and low-quality bases were trimmed off using Cutadapt (v2.10; [8]). The quality-filtered reads were assembled using MEGAHIT (v1.1.2; [9]) with default parameters. Contigs shorter than 1 kb were removed using Seqtk (v1.3; [10]) to ensure high-quality assembly. The quality-filtered reads were mapped back to the assembled contigs using Minimap2 (v2.17; [11]). The resulting alignment files were processed and converted from SAM to BAM format, followed by sorting and indexing using SAMtools (v1.11; [12]). Genome binning was performed using MetaBAT2 (v2.15; [13]), applying default parameters to extract metagenome-assembled genomes (MAGs). Together, this approach yielded 7 complete or high-quality genomes.

4. Additional methods of comparative genomics

To investigate the distribution and abundance patterns of antioxidant and desiccation-resistant genes identified in the genomes of the analysed methanogens, a gene copy number matrix was constructed (Table S7). Bray-Curtis dissimilarities were calculated from this matrix using the ‘vegan’ package (v2.6-4) in R [14], followed by Principal Coordinates Analysis using the base function ‘cmdscale’ [15]. The first two ordination axes were visualized with ‘ggplot2’.

For global comparisons of methanogen presence and abundance in desert soils, raw metagenome datasets (excluding amplicon studies) were downloaded from NCBI (Table S4). Searches were conducted for Bioprojects with the keywords ‘dry’, ‘oxic’, ‘desert’, ‘Arctic’, ‘Antarctic’, and ‘soil crust’. Using this approach, 24 Bioprojects (799 datasets) were identified and downloaded. We then used SingleM (v0.17.0) to profile the general taxonomic composition and identify the relative abundance of distinct methanogen taxa from the raw read data [16].

5. Pangenome construction and annotation using Anvi’o

Non-circular genomes were first filtered to retain contigs ≥ 2,500 bp using ‘anvi-script-reformat-fasta’ to ensure high-quality genomes. Each genome was then converted into an Anvi’o contigs database using ‘anvi-gen-contigs-database’. To facilitate functional and taxonomic annotation, we integrated the NCBI Clusters of Orthologous Groups (COGs) database and the single-copy gene (SCG) taxonomy database. Identifications for rRNA genes, single-copy core genes, and functional categories were performed using ‘anvi-run-hmms’, ‘anvi-run-ncbi-cogs’, and ‘anvi-scan-trnas’. We confirmed genome completeness (≥ 90%) and redundancy (≤ 10%) using ‘anvi-estimate-genome-completeness’ to ensure the reliability of comparison. All genomes of each methanogen class were then separately merged into single datasets using ‘anvi-gen-genomes-storage’. Afterwards, the pangenome analysis was conducted with ‘anvi-pan-genome’, which clusters orthologous genes into groups and classifies them either as core genes (shared across all genomes), accessory genes (present in some but not all), and unique genes (found in only one genome). Pangenome clustering results were visualized using ‘anvi-display-pan’, allowing for the comparison of gene presence or absence across multiple genomes.

**References**

1.	DSMZ. 141: *Methanogenium* medium (H_2_/CO_2_). 2024.

2.	DSMZ. 318: *Methanosarcina thermophila* (BCYT) medium. 2022.

3.	DSMZ. 1523: Modified *Methanobacterium* medium. 2023.

4.	Leibniz Institute DSMZ: *Methanocella paludicola* (DSM 17711). DSMZ.de. https://www.dsmz.de/collection/catalogue/details/culture/DSM-17711.

5.	Hori T, Aoyagi T, Itoh H. et al. Isolation of microorganisms involved in reduction of crystalline iron(III) oxides in natural environments. *Front Microbiol* 2015;**6**. https://doi.org/10.3389/fmicb.2015.00386

6.	DSMZ. Se/W solution. https://bacmedia.dsmz.de/download/solution/4034/pdf

7.	Angel R, Petrova E, Lara-Rodriguez A. Total nucleic acids extraction from soil. protocols.io. https://www.protocols.io/view/total-nucleic-acids-extraction-from-soil-bwxcpfiw. (27 July 2021, date last accessed).

8.	Martin M. Cutadapt removes adapter sequences from high-throughput sequencing reads. *EMBnet J* 2011;**17**:10–12. https://doi.org/10.14806/ej.17.1.200

9.	Li D, Liu C, Luo R. et al. MEGAHIT: an ultra-fast single-node solution for large and complex metagenomics assembly via succinct de Bruijn graph. *Bioinformatics* 2015;**31**:1674–1676. https://doi.org/10.1093/bioinformatics/btv033

10.	Li H. Toolkit for processing sequences in FASTA/Q formats. github.io. https://github.com/lh3/seqtk. .

11.	Li H. Minimap2: pairwise alignment for nucleotide sequences. *Bioinformatics* 2018;**34**:3094–3100. https://doi.org/10.1093/bioinformatics/bty191

12.	Danecek P, Bonfield JK, Liddle J. et al. Twelve years of SAMtools and BCFtools. *GigaScience* 2021;**10**:giab008. https://doi.org/10.1093/gigascience/giab008

13.	Kang DD, Li F, Kirton E. et al. MetaBAT 2: an adaptive binning algorithm for robust and efficient genome reconstruction from metagenome assemblies. *PeerJ* 2019;**7**:e7359. https://doi.org/10.7717/peerj.7359

14.	Oksanen J, Simpson G, Blanchet FG. et al. vegan community ecology package version 2.6-2 April 2022. 2022.

15.	R Core Team. R: A language and environment for statistical computing. 2021. R Foundation for Statistical Computing, Vienna, Austria, 2021.

16.	Woodcroft BJ, Aroney STN, Zhao R. et al. SingleM and Sandpiper: Robust microbial taxonomic profiles from metagenomic data. 2024. *bioRxiv* 2024.01.30.578060

**Supplementary Figures**

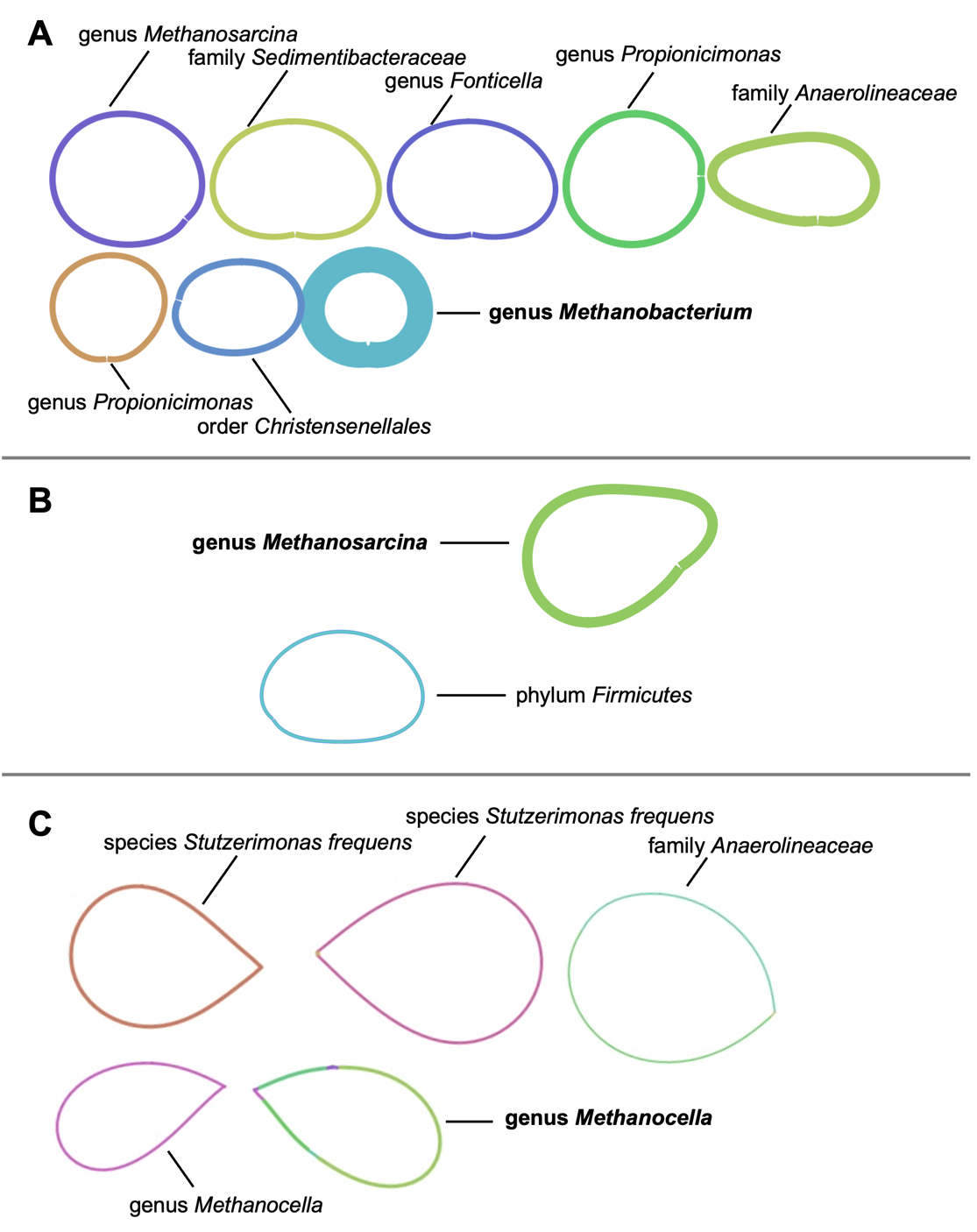

Figure S1. Closed genomes assembled with Flye and visualized using Bandage. The target genomes are indicated in bold font: (A) Methanobacterium sp., (B) Methanosarcina sp., and (C) Methanocella sp.

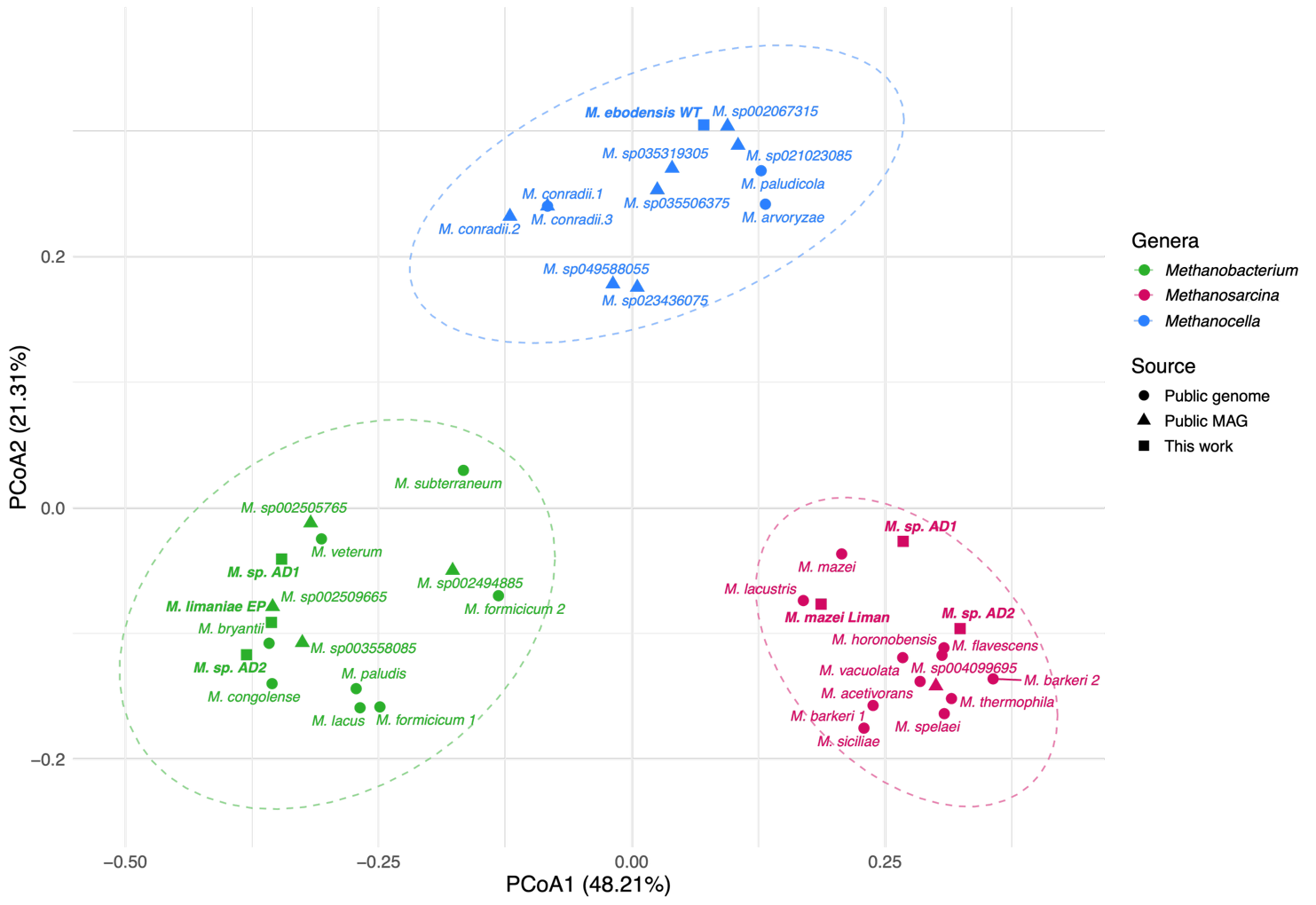

Figure S2. Principal Coordinates Analysis of methanogen gene abundance and diversity related to oxidative stress- and desiccation-resistance. Shapes represent genome source: circles are public genomes, triangles are public MAGs, and squares are genomes in this study. Colors distinguish genera: green for Methanobacterium, red for Methanosarcina, and blue for Methanocella. Dotted lines denote 95% confidence intervals.

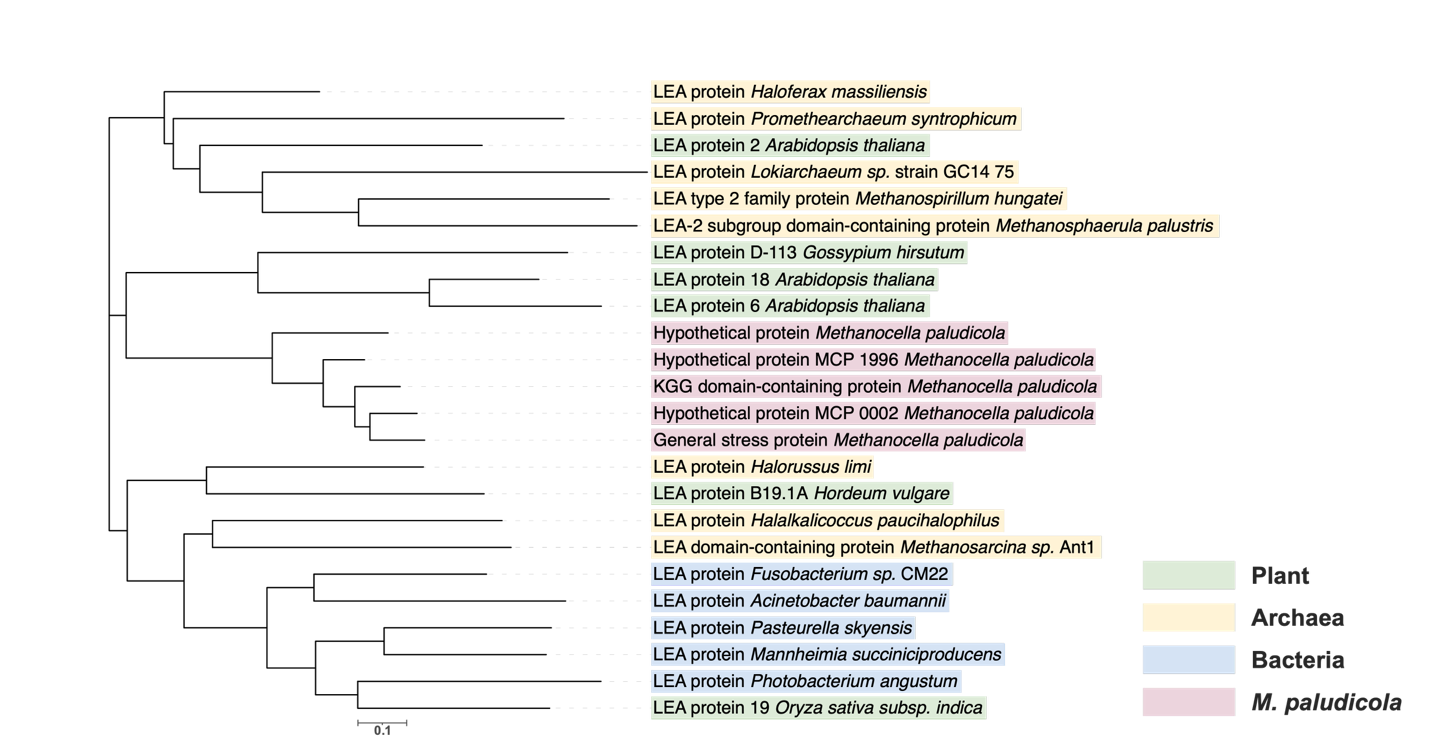

Figure S3. Neighbor-joining phylogenetic tree of LEA proteins. Proteins annotated as LEA proteins by Campos et al. are highlighted in pink (here named according to the results of tBLASTn). Verified LEA proteins found in plant, archaea, and bacteria are highlighted in green, yellow, and blue, respectively.

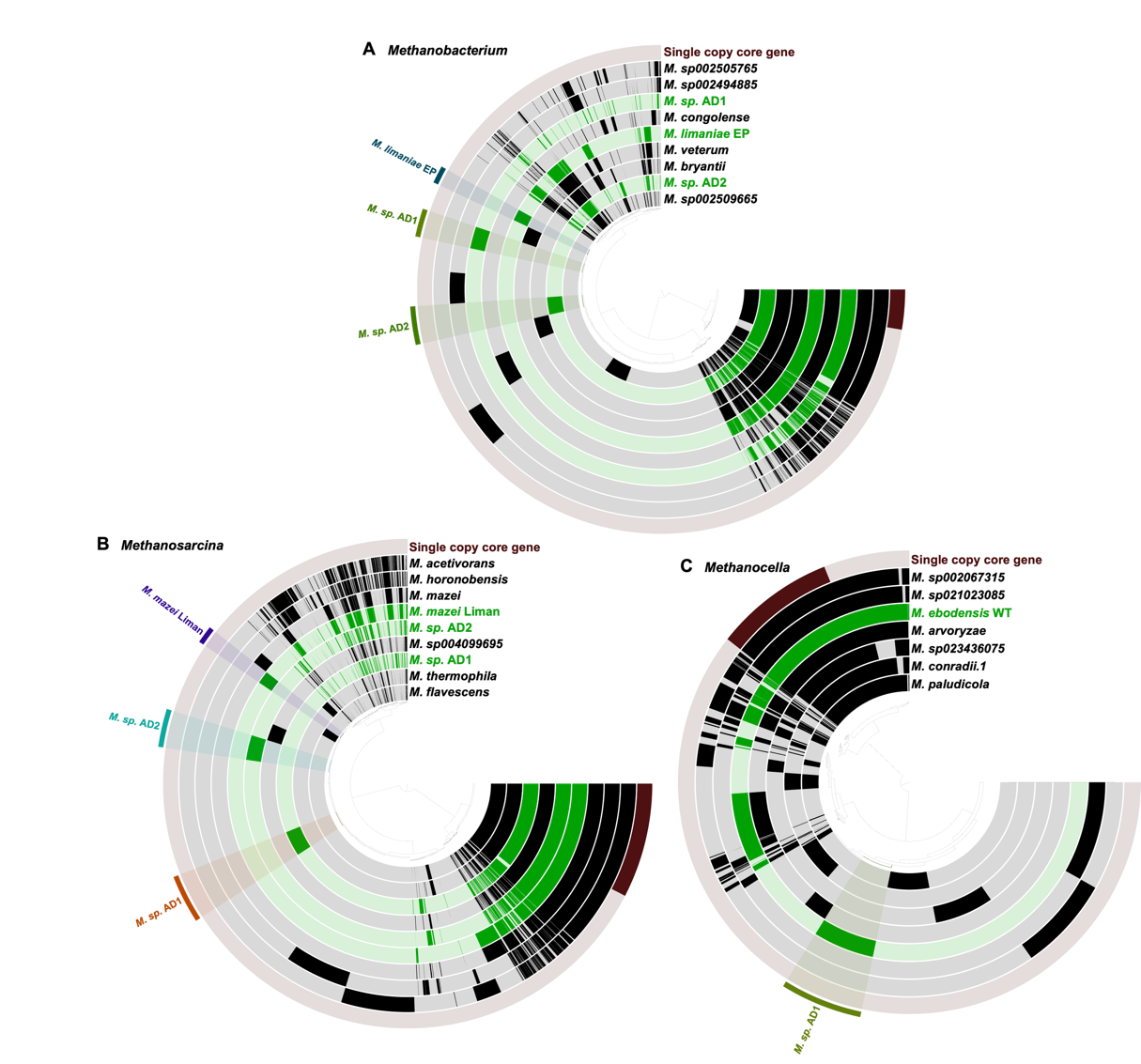

Figure S4. Pangenome analyses of the three methanogen genera: Methanobacterium (A), Methanosarcina (B), and Methanocella (C). Closely related high quality genomes to genomes generated in this study (highlighted in green) were selected. Unique gene clusters in genomes of this study are highlighted and labled.

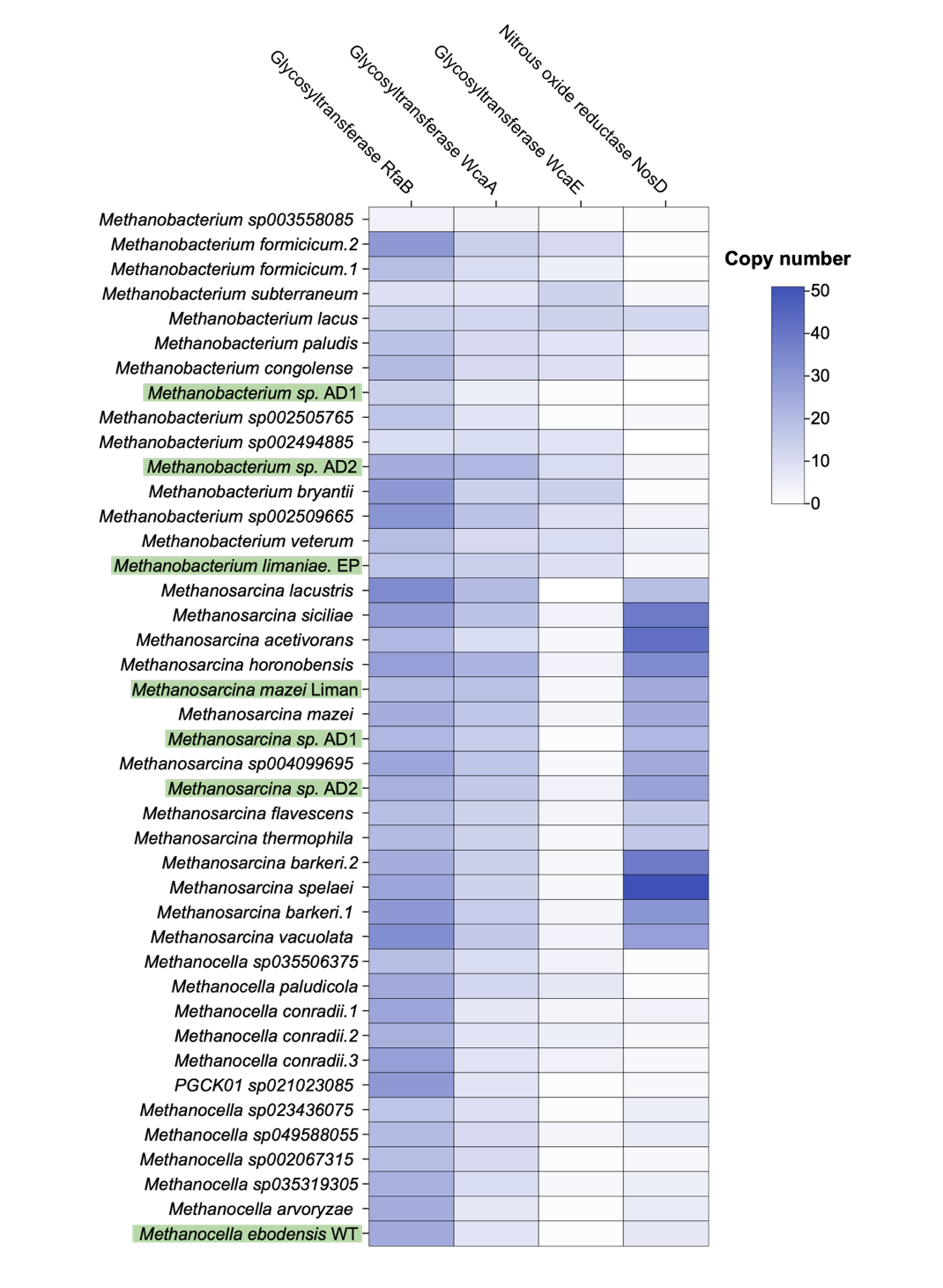

Figure S5. Gene copy number heatmap of glycosyltransferase rfaB, glycosyltransferase wcaA, glycosyltransferase wcaE, and nitrous oxide reductase nosD found by anvi’o annotation in selected genomes. Genomes sequenced in this study are highlighted in green.

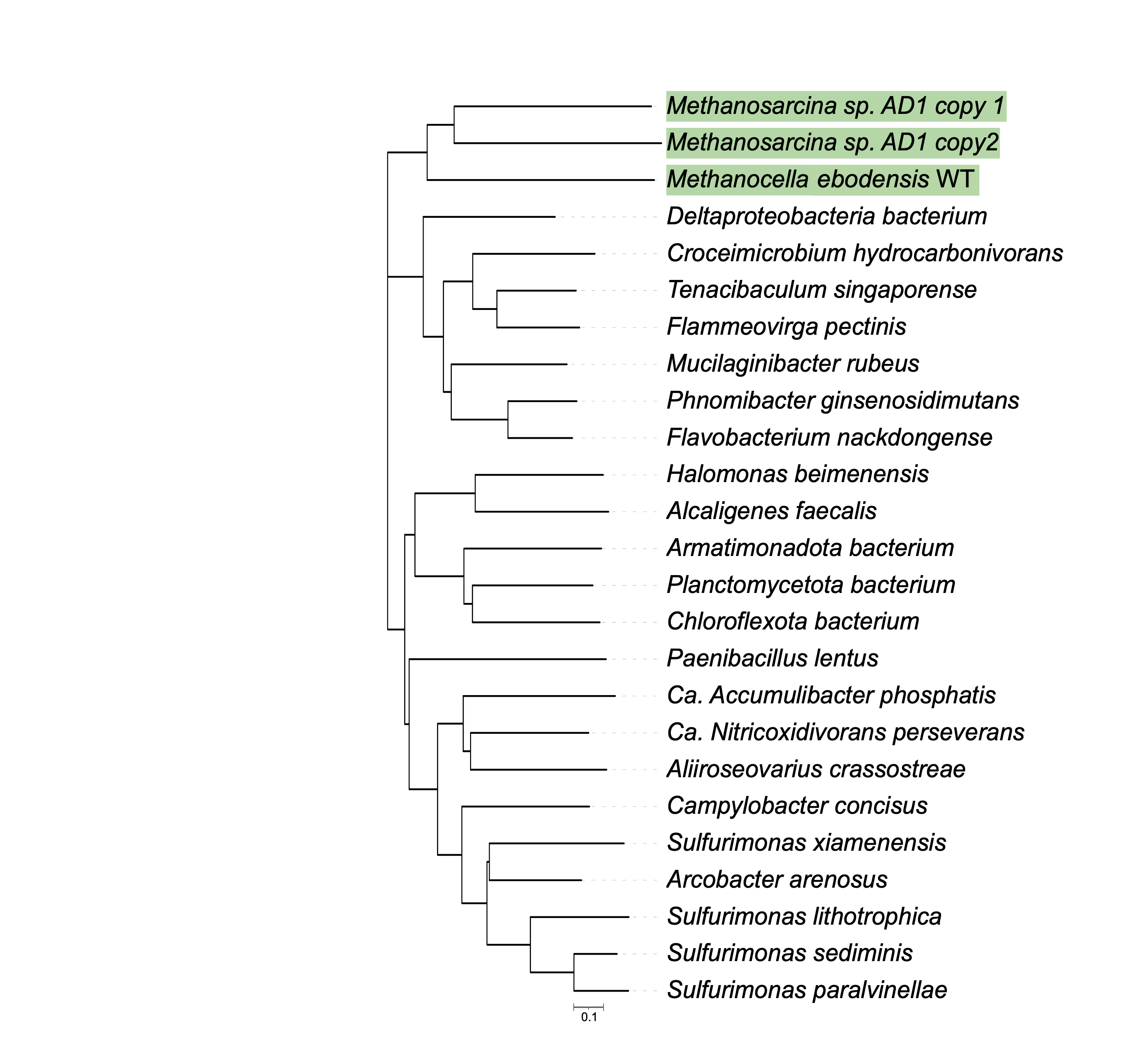

Figure S6. Neighbor-joining phylogenetic protein tree of the nitrous oxide reductase subunit NosD found in different microorganisms. Three NosD proteins found in the unique clusters of genomes from this study are highlighted in green.

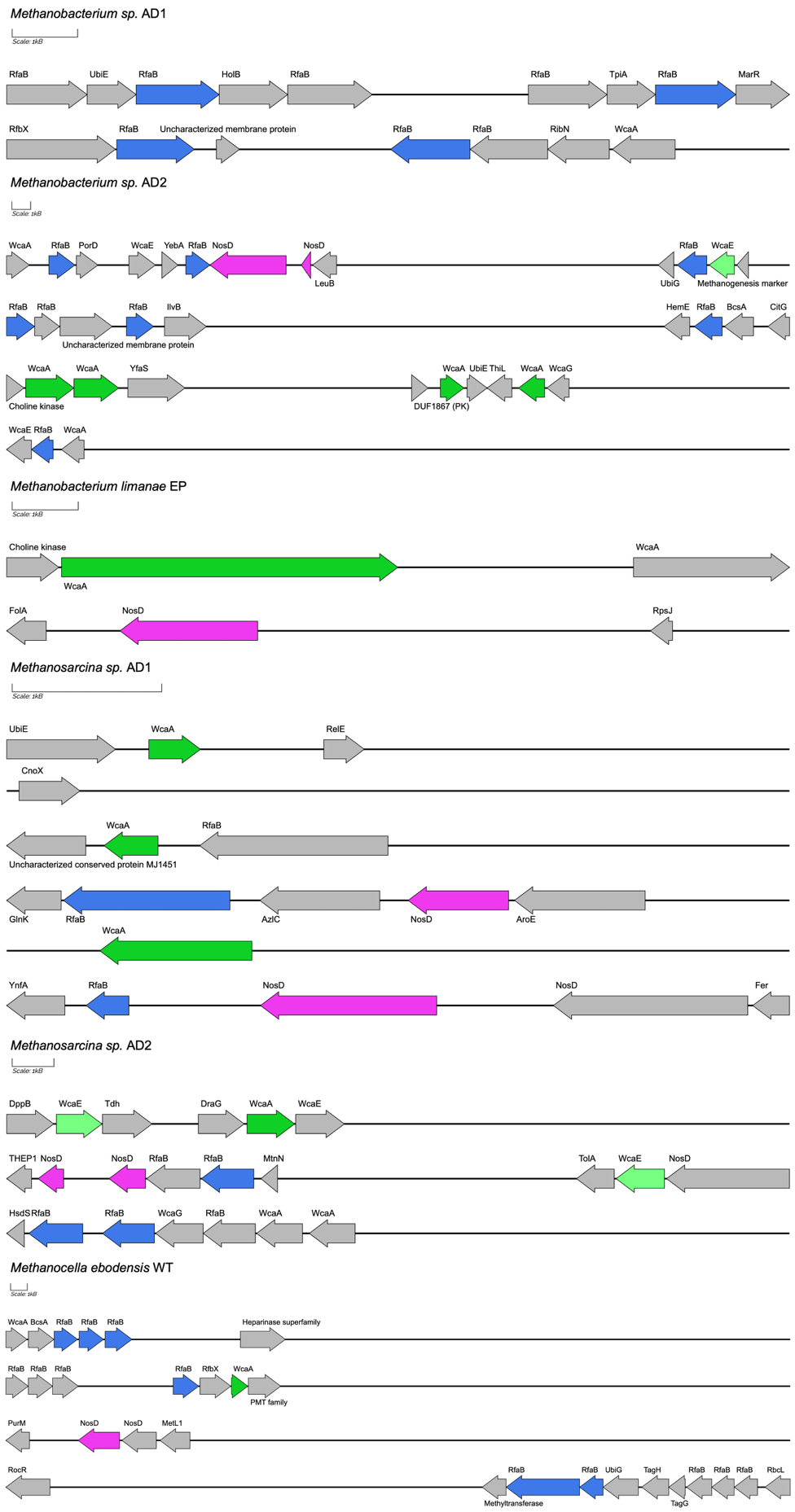

Figure S7. Synteny and genomic context of rfaB, wcaA, wcaE, and nosD genes in the genomes. Genes identified as unique are shown in color (blue, RfaB; green, WcaA and WcaE; and pink, NosD). These 4 genes were not among the unique genes of Methanosarcina mazei Liman.
